# Defining the emergence of myeloid-derived suppressor cells in breast cancer using single-cell transcriptomics

**DOI:** 10.1101/702860

**Authors:** Hamad Alshetaiwi, Nicholas Pervolarakis, Laura Lynn McIntyre, Dennis Ma, Quy Nguyen, Jan Akara Rath, Kevin Nee, Grace Hernandez, Katrina Evans, Leona Torosian, Anushka Silva, Craig Walsh, Kai Kessenbrock

## Abstract

Myeloid-derived suppressor cells (MDSCs) are innate immune cells that acquire the capacity to suppress adaptive immune responses during cancer. It remains elusive how MDSCs differ from their normal myeloid counterparts, which limits our ability to specifically detect and therapeutically target MDSCs during cancer. Here, we used single-cell RNAseq to compare MDSC-containing splenic myeloid cells from breast tumor-bearing mice to wildtype controls. Our computational analysis of 14,646 single-cell transcriptomes reveals that MDSCs emerge through a previously unrealized aberrant neutrophil maturation trajectory in the spleen giving rise to a unique chemokine-responsive, immunosuppressive cell state that strongly differs from normal myeloid cells. We establish the first MDSC-specific gene signature and identify novel surface markers for improved detection and enrichment of MDSCs in murine and human samples. Our study provides the first single-cell transcriptional map defining the development of MDSCs, which will ultimately enable us to specifically target these cells in cancer patients.

**One Sentence Summary:** We used single cell transcriptomics to identify the unique molecular features distinguishing myeloid-derived suppressor cells (MDSCs) from their normal, myeloid counterparts, which enabled us to reveal distinct transitory gene expression changes during their maturation in the spleen, and to identify novel cell surface markers for improved detection and isolation of MDSCs.

## Introduction

Breast cancer is one of the most prevalent types of cancer with over 260,000 new cases and over 40,000 deaths in 2018 in the US^1^. During tumor development, breast cancer cells secrete various cytokines such as granulocyte-macrophage colony-stimulating factor (GM-CSF), which exert systemic effects on hematopoiesis and myeloid cell differentiation promoting the development of myeloid-derived suppressor cells (MDSCs)^2, 3^. These MDSCs are a heterogeneous population of neutrophil- and monocyte-like myeloid cells, which are increasingly recognized as key mediators of immune suppression in various types of cancer^3, 4^. In cancer patients, increased numbers of MDSCs in circulation correlate with advanced clinical stages, increased metastatic progression and immune suppression^5^. MDSCs can mediate immune suppression through multiple mechanisms including the production of reactive oxygen species (ROS) and depletion of key amino acids required for T cell proliferation through expression of arginase (Arg) and indoleamine 2,3-dioxygenase (IDO)^6–8^. In addition, MDSCs produce a range of immunosuppressive and cancer-promoting cytokines including IL-10 and TGF-β^9^. Besides their immune-suppressive function, MDSCs may also actively shape the tumor microenvironment through complex crosstalk with breast cancer cells and surrounding stroma, resulting in increased angiogenesis, tumor invasion, and metastasis^8,10, 11^.

The unique molecular features of MDSCs are currently unclear and it remains elusive whether MDSCs represent a unique subpopulation of myeloid cells that differ from their normal, healthy counterparts. This limits our ability to determine specific MDSC functions as opposed to bulk-level changes in neutrophils or monocytes during cancer. In mice, MDSCs are defined through the expression of CD11b^+^Gr1^+^ and can be further classified into CD11b^+^Ly6C^low^Ly6G^+^ granulocytic MDSCs (G-MDSCs) and CD11b^+^Ly6C^+^Ly6G^-^ monocytic MDSCs (M-MDSCs)^12^. In humans, G-MDSCs are defined as CD11b^+^CD14^-^CD15^+^ or CD11b^+^CD14^-^CD66b^+^and M-MDSCs as CD11b^+^CD14^+^HLA-DR^-/low^CD15^-^ followed by additional functional characteristics such as T cells suppression and ROS assays^12^. However, these markers overlap with those defining healthy neutrophils and monocytes, which makes it challenging to distinguish MDSCs from normal cells to advance our understanding of MDSCs biology and ultimately, to establish novel therapeutic avenues to interfere with their tumor-promoting and immune suppressive roles.

Here, we used single-cell RNA sequencing (scRNAseq) to delineate the unique molecular features of MDSCs in the MMTV-PyMT mouse model of breast cancer. Our computational analysis of 14,646 single cell transcriptomes revealed a unique MDSC gene signature, which is largely shared between G-MDSCs and M-MDSCs, but which strongly differs from their normal myeloid counterparts. Focusing on G-MDSCs, our pseudotemporal analysis delineates the emergence of MDSCs as an aberrant differentiation state that forms a separate branch during the transition of neutrophil progenitors into mature neutrophils. Further interrogation of the distinct MDSC gene expression signature identified several novel surface markers (e.g. CD84, JAML) for faithful MDSC detection and prospective enrichment. Taken together, our study provides the first single-cell level molecular census defining novel specific gene signatures and markers for MDSCs that were previously unrealized in bulk-level expression analyses, which may form the foundation to ultimately therapeutically interfere with MDSC function in cancer patients.

## Results

### Spleen is the predominant organ of MDSC generation in tumor-bearing mice

Mice expressing the polyomavirus middle T antigen (PyMT) driven by the mouse mammary tumor virus (MMTV) promoter^13^ develop breast tumors that closely resembles human pathogenesis^14^ and give rise to MDSCs during tumor progression^2^. Here, we used the MMTV-PyMT transgenic mouse model of breast cancer to explore the role of MDSCs during breast cancer progression. We first sought to confirm the most reliable organ site of MDSC accumulation for further molecular studies of this cell population. In accordance with previous reports in other murine models of cancer^2^, we observed that later stages of cancer progression were associated with an expansion of CD11b^+^Gr1^+^ myeloid cells in bone marrow, blood, spleens, lungs, brains and primary tumors (**fig. S1A-C**), and an enlargement of the spleen of tumor-bearing PyMT mice compared to wildtype (WT) controls (**fig. S1D-E**). To functionally confirm whether MDSCs are present in expanded populations of CD11b^+^Gr1^+^ cells in tumor-bearing mice, we isolated CD11b^+^Gr1^+^ cells from various organs of tumor-bearing and control mice by fluorescence-activated cell sorting (FACS) and co-cultured these with isolated T cells to measure suppression of T cell proliferation induced by CD3/CD28 co-stimulation^2^, and reactive oxygen species (ROS) formation as a read-out for MDSC function^12^ (**fig. S2A**). We found that CD11b^+^Gr1^+^ cells sorted from spleens of tumor-bearing mice significantly suppressed CD4^+^ and CD8^+^ T cell proliferation (**fig. S2B-C**), whereas CD11b^+^Gr1^+^ cells from control spleens showed no measurable effect on T cell proliferation. Of note, CD11b^+^Gr1^+^ cells sorted from bone marrow (**fig. S2D-E**) and lungs (**fig. S1F-G**) of tumor-bearing mice demonstrated only nonsignificant suppression of T cell proliferation. These findings were further corroborated by ROS production assays as measured by flow cytometry using 2ʹ,7ʹ-Dichlorofluorescin diacetate (H_2_DCFDA), which showed that only spleen-derived CD11b^+^Gr1^+^ cells from tumor-bearing mice exhibited significant oxidative burst formation as a hallmark for MDSCs (**fig. S2F-G**). Together, these results establish the spleen as the major site of MDSC emergence during breast tumor formation in PyMT mice.

### Single-cell transcriptomics reveal MDSCs as distinct clusters within neutrophilic and monocytic lineages

In order to determine how MDSCs differ from their normal myeloid counterparts on a cellular and molecular level, we used scRNAseq to compare the molecular differences of spleen-derived myeloid cells in tumor-bearing mice against the respective cell population from WT mice on an individual cell basis. We utilized a scalable droplet-mediated scRNAseq platform (10X Genomics Chromium) to profile FACS-purified live (Sytox-negative) CD45^+^CD11b^+^Gr1^+^ myeloid cells from the spleens of tumor-bearing PyMT and control WT mice (**Fig. 1A**). We profiled two samples from tumor-bearing PyMT mice (9,155 cells) and WT control mice (5,491 cells), respectively, for a total of 14,646 cells that were sequenced at an average depth of ∼50,000 reads per cell. The two libraries were aggregated and aligned together using the CellRanger pipeline (10X Genomics) to compensate for minor differences in library complexity. After quality control filtering to remove cells with low gene detection (<500genes) and high mitochondrial gene content (>8%), we performed clustering and cell type identification analysis of combined PyMT and WT datasets using Seurat^15^ (**fig. S3A**). Using the canonical correlation analysis (CCA) method^16^, we identified the main cell types based on expression of hallmark genes for myeloid subsets (**Fig. 1B**), and determined their marker genes (**fig. S3C**; **table S1**). Neutrophils formed the largest population encompassing numerous distinct clusters (C0, C2, C4, C5, C7 and C8) characterized by high levels of *Ly6g* and *Cxcr2* expression (**Fig. 1B-C**). Monocytes were less abundant and less diverse forming one cluster (C1) that was marked by expression of *Csf1r* and *Ccr2* (**Fig. 1B-C**). We also detected two minor cell types: T cells (C9) expressing *Cd3*, *Cd4*, *Cd8*; and B cells (C6 and C3) expressing *Cd19*, *Cd22*, *Cd79a* (**Fig. 1B-C**).

**Fig. 1.**
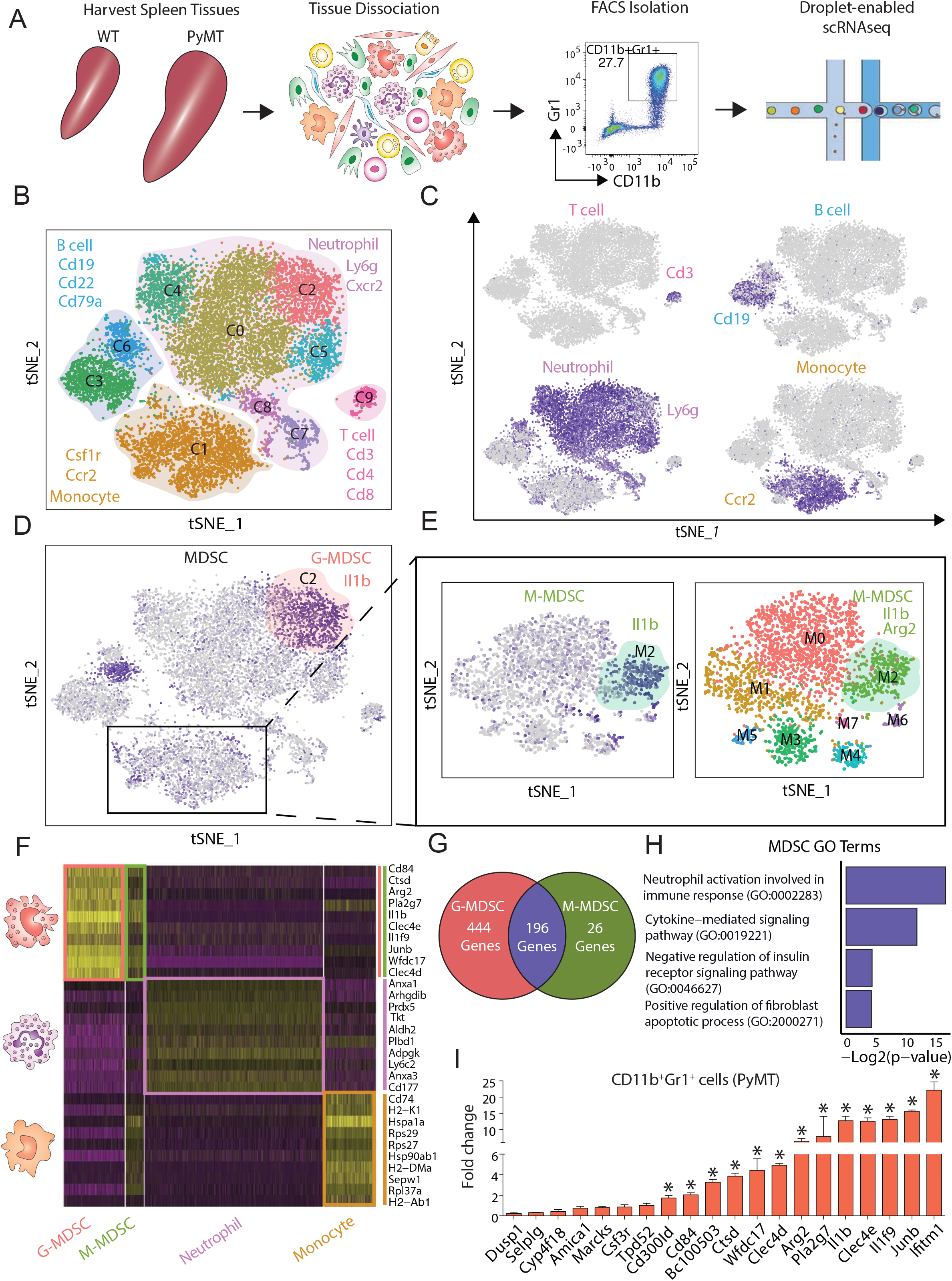
Identifying MDSC-specific gene expression signatures using scRNAseq. (**A**) Approach overview for single-cell analysis of and (sytox blue-negative) CD45^+^CD11b^+^Gr1^+^ cells were sorted from the spleen of control WT and tumor-bearing PyMT’s mice by FACS following droplet-enabled scRNAseq. (**B-C)** Combined Seurat analysis of in total 14,646 cells from control and PyMT mice shown in tSNE projection results in various distinct clusters of splenic CD11b^+^Gr1^+^ cells. Main cell types (T cells, B cells, neutrophils, monocytes) are outlined based on hallmark gene expression. (**C)** Feature plots of characteristic markers of the four main cell types showing expression levels with low expression in grey to high expression in dark blue. (**D**) G-MDSCs were identified in cluster C1 by expression marker genes (*Arg2* & *Il1β*) from the PyMT sample. (**E**) Subset analysis of monocytes cluster identified M-MDSCs. Eight total clusters were found; cluster M2 was identified as M-MDSCs (positive for *Arg2* & *Il1β*). (**F**) Heatmap displaying the scaled expression patterns of top marker genes within each G-MDSCs and M-MDSCs clusters compared to normal neutrophil and monocyte clusters from WT mice respectively; yellow = high expression; purple = low expression. (**G**) Venn diagram showing the number of statistically significant marker genes and overlap between G-MDSC and M-MDSC. (**H**) Gene ontology (GO) term analysis using Enrichr of curated MDSC signature. (**I**) Validation using qPCR of selected upregulated MDSC genes, statistical analysis unpaired t-test (Mean ± SEM of n = 3) *\*P<* 0.05.

Further interrogation of neutrophil heterogeneity revealed that cluster C0 was marked by high levels of genes associated with a mature neutrophil state such as *Camp*^17^ and high *Ly6g* expression^18^; cluster C2 was strongly enriched in tumor-bearing PyMT mice (**fig. S3E**) and displayed high expression MDSC-related genes such as *Il1β* and *Arg2*, two major immunosuppressive factors previously used to define MDSCs in cancer models^4, 19^ (**Fig. 1D**); clusters C4 and C5 displayed overlapping marker gene expression including genes such as *Cebpe* and *Retnig*; clusters C7 and C8 exhibited high expression of cell cycle genes such as *Tuba1b* and *Cdc20* indicating the existence of a proliferative pool of neutrophils in the spleen.

We next focused on the monocyte-restricted cluster C1, which showed diffuse expression of MDSC genes *Arg2* and *Il1b* in the combined analysis (**Fig. 1D**) suggesting that M-MDSC were present but not clustering distinctly from monocytes due to the more substantial differences between different cell types in the combined analysis. Therefore, we performed a monocyte-only clustering analysis to identify several distinct states (clusters M0-M7) including a distinct M-MDSC population in cluster M2, which was strongly enriched in tumor-bearing PyMT mice (**Fig. 1E**; **fig. S3B, D, F**; **table S2**). These analyses formed the basis for a detailed molecular definition of G- and M-MDSCs as described below. Our dataset represents the first single-cell level transcriptome analysis of MDSCs, which revealed that G- and M-MDSCs form distinct clusters that are unique from their normal myeloid counterparts.

### G- and M-MDSCs share a conserved immune cell activation program that strongly differs from normal myeloid cells

We next utilized our scRNAseq dataset to reveal the unique molecular features of MDSCs and to unravel the distinct biological programs that define the MDSC state. We performed differential expression analysis in Seurat to determine how G- and M-MDSCs from tumor-bearing mice differ from their normal counterparts, namely neutrophils and monocytes in WT mice (**Fig. 1F**). Our analysis revealed 642 differentially expressed genes in G-MDSCs compared to normal neutrophils (**table S3**), and 223 differentially expressed genes in M-MDSCs compared to normal monocytes (**table S4**) demonstrating that MDSCs differ substantially from their normal myeloid counterparts. Interestingly, there was substantial overlap between gene signatures for G- and M-MDSCs (196 genes, **Fig. 1G**; **table S5**) indicating that this immune-suppressive cell state can be acquired by both monocytes and neutrophils independently. Shared markers included genes involved in immune suppression such as *Il1b*, *Arg2*, *Cd84* and *Wfdc*^20^. Interferon-induced transmembrane protein 1 (*Ifitm1*), which has been reported to be involved in progression of colorectal cancer^21, 22^ and inflammatory breast cancer cells^23^ was upregulated in MDSCs. Additional MDSC markers included myeloid associated immunoglobulin like receptor family (*Cd300ld*), C-type lectin domain family 4-member E and D (*Clec4e* and *Clec4d*), Interleukin 1f9 (*Il1f9*), AP-1 transcription factor subunit (*Junb*), Cathepsin D (*Ctsd*), phospholipase A2 group VII (*Pla2g7*) and cystatin domain containing 5 (*Bc100530*).

We next performed gene ontology (GO) term analysis (**Fig. 1H**) using Enrichr (*GO Biological Process 2018*)^24^. The top GO terms included ‘*neutrophil activation and involved in immune response’* genes including the genes encoding complement C5a receptor 1 (*C5ar1*), S100 Calcium binding protein A11(*S100a11*), *Clec4d*, chemokine receptor 2 (*Cxcr2),* and Annexin A2 (*Anxa2*). Interestingly, several of these factors promote recruitment of neutrophils and MDSCs as reported for *C5ar1*^25, 26^, and *S100a8/9*^27, 28^. In addition, *Anxa2* has been reported to modulate ROS production and inflammatory responses^29^, which is a hallmark of MDSCs. Another significant GO term *‘Cytokine-mediated signaling pathway’* included genes such as *Il1β, Ifitm1*, *Junb*, and *Myd88*. In particular, *Myd88* has been reported to promote expansion of immature Gr1+ cells and may be involved in mediating T cell suppressing cell states^30^. Additionally, this pathway included genes associated with MDSCs accumulation and trafficking such as *Cxcr2*^31, 32^, *Csf3r*^33^ and *Ccr1*^34^, suggesting that MDSCs may be able to responsive to recruitment signals from sites of inflammation such as the primary tumor or metastatic sites. Moreover*, ‘Negative regulation of insulin receptor signaling pathway’* genes such as; *Il1β* ^35^, and suppresser of cytokine signaling 3 (Socs3) were prominent in MDSCs. Socs3 has been reported to regulate granulocyte colony stimulating factor (G-CSF), and signal transducer and activator of transcription 3 (STAT3) activation^36^ (**table S6**).

Next, we sought to orthogonally validate the MDSC genes signature. To this end, CD11b^+^Gr1^+^ cells from spleens of WT and tumor-bearing PyMT mice were isolated by FACS and subjected to quantitative PCR (qPCR). Our scRNAseq results were broadly confirmed in this targeted approach, since a large proportion of MDSC signature genes were significantly upregulated in CD11b^+^Gr1^+^ cells from PyMT compared to WT (**Fig. 1I**). Taken together, these analyses firstly revealed that G- and M-MDSCs share aconserved gene signature that strongly differs from their normal myeloid counterparts. This shared MDSC marker gene list differs significantly from previous transcriptome-level analyses of MDSCs, indicating that bulk-level changes in these myeloid cell populations mask the specific programs underlying MDSC cell function.

### MDSC gene signature is highly expressed in human breast cancer-associated neutrophils

To determine whether this MDSC gene signature is generalizable and translatable into the human context, we explored a recently published scRNAseq immune cell map including T cells, B cells, monocytes, neutrophils from primary tumor samples of breast cancer patients^37^. We performed clustering of this dataset in Seurat to reproduce cell type labels (**Fig. 2A**), and then carried out an unbiased gene signature scoring of all cell types, which revealed that specifically neutrophils and monocytes in the tumor microenvironment express high levels of MDSC signature genes (**Fig. 2B**). To assess whether there are distinct subsets of neutrophils with particularly high MDSC signatures, we also analyzed neutrophils separately using unbiased clustering yielding four distinct states (**Fig. 2C-D**). Interestingly, cluster 0 showed MDSC-related marker genes *S100A9* and *CCR2*, suggesting that this subset of neutrophils represents G-MDSCs in the tumor microenvironment of breast cancer patients. Gene scoring analysis of using our MDSC gene signature in these neutrophil subclusters indeed showed by far the highest scores in cells from cluster 0 (**Fig. 2E**). Together, these analyses confirmed that our MDSC gene signature derived from a murine breast cancer model is translatable into human disease indicating that the MDSC state is largely conserved between mice and human.

**Fig. 2.**
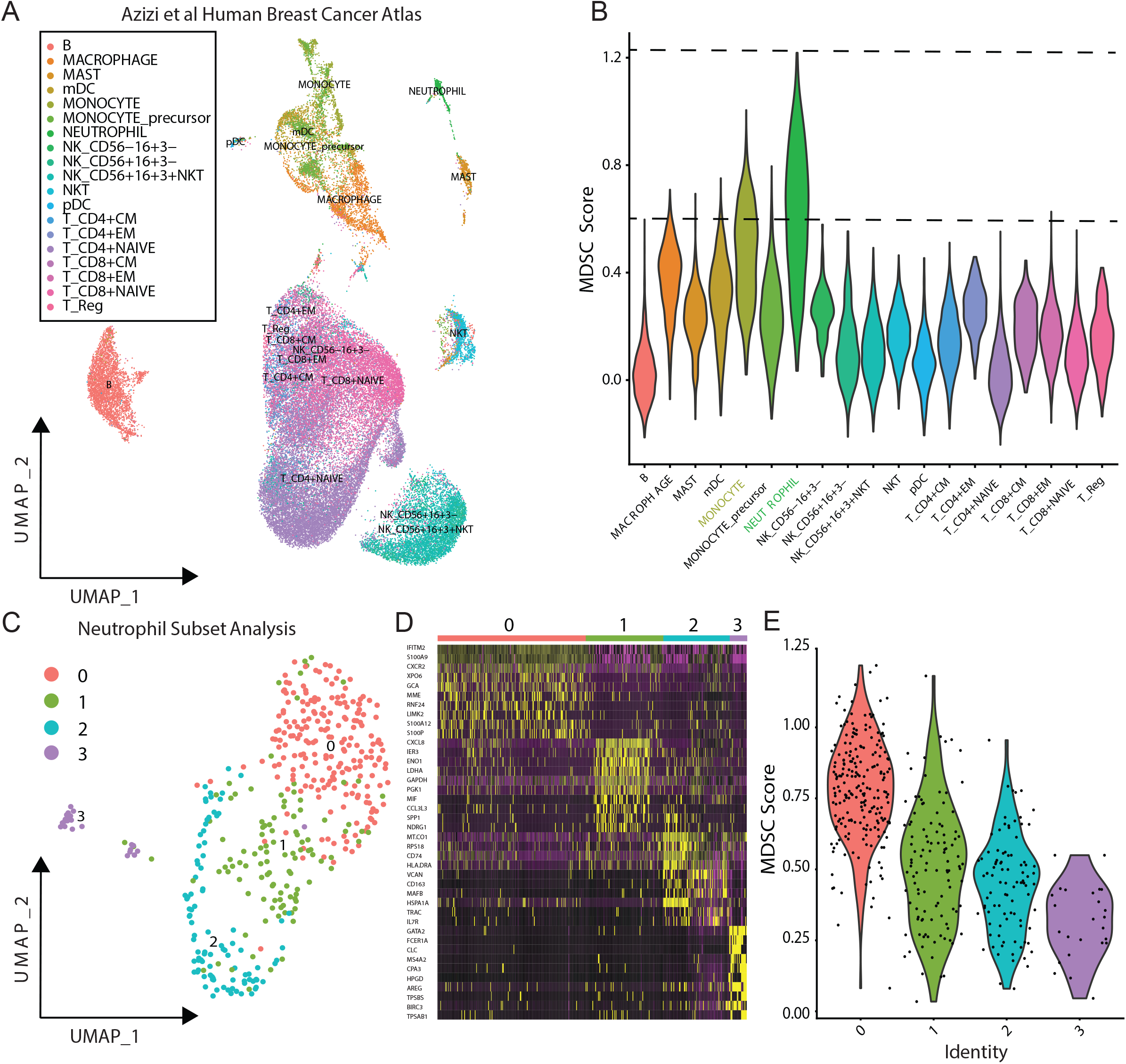
MDSC signature marks subset of neutrophils in the tumor microenvironment of human breast cancer patients. (**A**) Seurat analysis of scRNAseq dataset comprising various immune cell populations in primary human breast tumor samples^37^ projected in UMAP with cell type labels as indicated in different colors. (**B**) Violin plot showing relative MDSC score of all cells in this dataset ordered by cell type showing highest scores in neutrophils. (**C**) Separate unbiased Seurat clustering analysis of neutrophil alone projected in UMAP yielded four distinct clusters of neutrophils in this dataset. (**D**) Heatmap showing top 10 marker genes for each neutrophil cluster. (**E**) Violin plots showing relative MDSC score ordered by neutrophil subcluster showing that cluster 0 exhibit highest expression of MDSC gene signature.

### Aberrant neutrophil differentiation in the spleen gives rise to MDSCs in cancer

To reconstruct the maturation process leading to MDSC generation in the spleen and to determine their differentiation state relative to normal progenitor and mature neutrophil populations, we next performed Monocle for unsupervised pseudotemporal ordering of our scRNAseq dataset^38^. We focused on the *Ly6g+* neutrophil subset (clusters C0, C2, C4, C5, C7 and C8 in **Fig. 1B-C**) because in contrast to M-MDSCs we recovered sufficient numbers of total neutrophils and G-MDSCs in this analysis to ensure interpretable result. We first generated a new Seurat-based clustering of this neutrophil subset and then performed Monocle using this newly defined set of marker genes (**fig. S4A**; **table S7**). This resulted in a three-branch trajectory with 5 distinct cell states (**Fig. 3A**). To interpret this trajectory, we compared our results to recent work using scRNAseq to define the signatures of the naïve haematopoietic stem, progenitor and differentiated cell states in the bone marrow of mice, which revealed that neutrophil progenitors are marked by the genes *Elane*, *Mpo* and *Prtn3*, while mature neutrophils expressed elevated levels of *Camp*, *Ltf* and *Lcn2*^17^. Integrating these markers together with the MDSC signature established in our work (**Fig. 1F**), we were able to annotate the five states. First identified were neutrophil progenitors (state 4; *Elane*-hi) that show increased proliferation (**fig. S4B-C**) and form the beginning of pseudotime. These progenitors then bifurcate into mature neutrophils (state 3; *Camp*-hi) on the one branch, and MDSCs (state 1; *Cd84*-hi) on the other branch as illustrated by gene plots over pseudotime (**Fig. 3B**), suggesting that MDSCs emerge from neutrophil progenitors via an alternative maturation process.

**Fig. 3.**
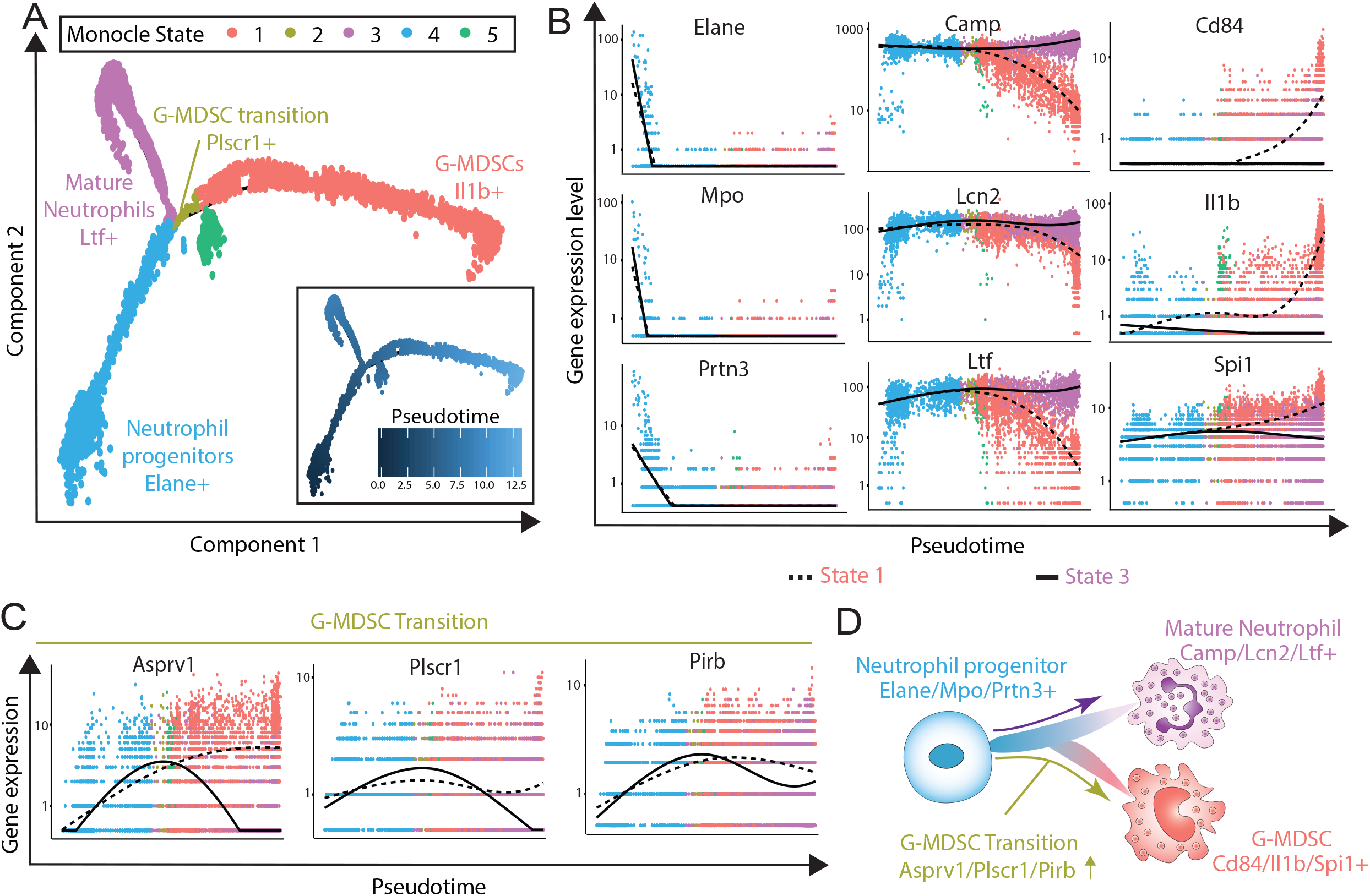
G-MDSCs emerge through aberrant differentiation trajectory during cancer. (**A**) Monocle analysis on the subset of Ly6g+ neutrophil clusters resulted in branched trajectory with 5 distinct Monocle states (color code for each state is indicated) which are named based on respective gene expression profile. (**B**) Pseudotime plot illustrating expression of selected marker genes over pseudotime with the branch ending in State 1 shown with the dotted line, and the branch ending with state 3 highlighted by the solid line. Neutrophil progenitors are characterized by high levels of *Elane, Mpo and Prtn3* (state 4), which bifurcate into mature neutrophils (state 3; *Camp*, *Ltf, Lcn2*) on the one branch, and MDSCs (state 1; *e.g. CD84*) on the other branch. **(C**) Early G-MDSC transition was marked by high expression of *Asprv1, Plscr1 and Pirb*. (**D**) Summary schematic indicates that G-MDSCs emerge from neutrophil progenitor cells via an aberrant form of neutrophil differentiation rather than from mature neutrophils that are reprogrammed into immunosuppressive cells.

Interestingly, Monocle detected two additional cell states (2 and 5) around the beginning of the MDSC branch: while state 5 was characterized by high ribosomal gene counts indicative of a translationally active cell state, state 2 represents the earliest phase of MDSC differentiation and was marked by high expression of *Asprv1*^39^, *Plscr1*^40^ and *Pirb*^41^ (**Fig. 3C**). Interestingly, it has been reported that neutrophils promote chronic inflammation using *Asprv1*^39^, suggesting the aspartic protease encoded by *Asprv1* may functionally contribute to the emergence of MDSCs in the spleen. Furthermore, paired immunoglobin-like receptor-b (*Pirb*) has been reported to regulate the suppressive function and fate of MDSC, indicating that *Pirb* is required for MDSC generation^41^. Taken together, these findings indicate that MDSCs emerge from neutrophil progenitor cells via an aberrant form of neutrophil differentiation in the spleen rather than from mature neutrophils that are reprogrammed into immunosuppressive cells (**Fig. 3D; table S8**). In addition, our work firstly identified an early, transitional MDSC state characterized by a number of genes showing elevated expression only around the branching point and during MDSC differentiation, but not during the normal progenitor or mature neutrophil trajectory. This may suggest that the transitional MDSC state could be targeted to block differentiation into MDSCs while not affecting normal neutrophil maturation and function.

### Identification of novel cell surface markers for MDSC detection and isolation

Our scRNAseq data revealed several previously unknown specific cell surface markers for MDSCs including CD84 and Amica1/Jaml. CD84 is a cell surface receptor of the signaling lymphocytic activation molecule (SLAM) family^42^ and is expressed on some immune cell types^43, 44^. Amica1/Jaml is a junctional adhesion molecule known to mediate the transmigration of neutrophils and monocytes by interacting with coxsackie-adenovirus receptor (CAR) expressed by epithelia^45^. We profiled CD84 and Jaml expression using FACS on CD11b^+^Gr1^+^ cells from different organs in tumor-bearing PyMT mice and WT mice. We first used FMO and isotype controls to determine specific marker expression (**fig. S4D-E**). Next, we characterized CD84 and Jaml expression in the CD11b^+^Gr1^+^ population from various organ preparations (bone marrow, lung, spleen, MFP or primary tumor) and compared control WT to tumor-bearing PyMT mice. Importantly, while CD11b^+^Gr1^+^ cells from bone marrow and lung were generally negative for CD84 (**Fig. 4A**) and Jaml (**Fig. 4D**), we found a significant number of CD11b^+^Gr1^+^ cells from the spleen and primary tumors of PyMT mice exhibited high expression of CD84 (**Fig. 4B**) and Jaml (**Fig. 4E**) compared to the respective WT controls. This is particularly apparent when cells from all organs are plotted side by side (**Fig. 4C&F**). This observation of high expression of CD84 and Jaml in spleen and primary tumors correlates with high MDSC capacity of CD11b^+^Gr1^+^ cells in these sites.

**Fig. 4.**
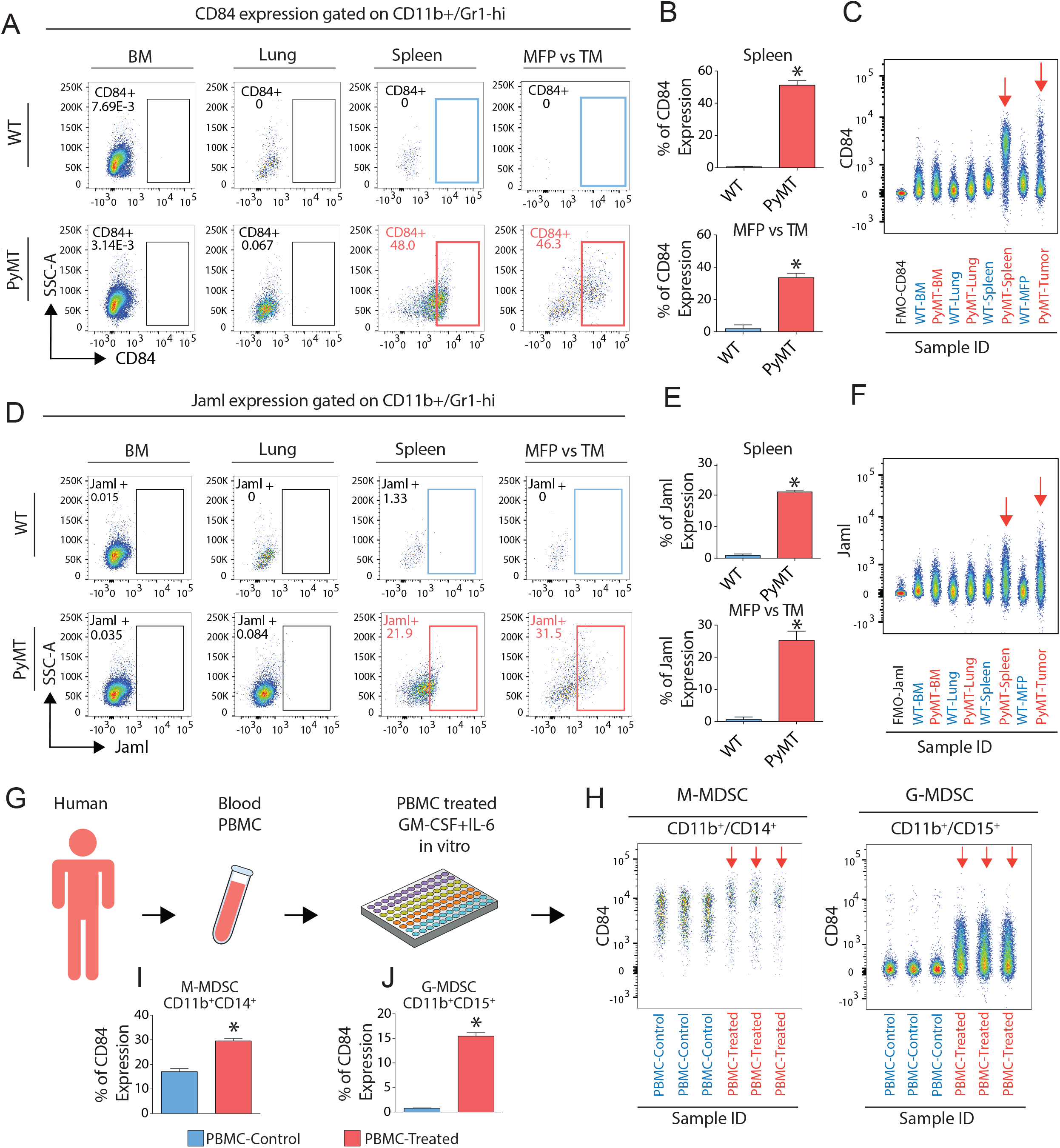
Identification of novel cell surface markers for MDSCs in breast cancer models. (**A**) CD84 expression profiling in WT and tumor-bearing PyMT showing that only spleen and primary tumor from PyMT exhibit significant expression. (**B**) Combined results and statistical analysis using unpaired t-test (Mean ± SEM of n = 10) *\*P<* 0.05. (**D**) Profiling Jaml expression in WT and PyMT showing only spleen and tumor from PyMT exhibit significant expression. (**E**) Combined results and statistical analysis unpaired t-test (Mean ± SEM of n = 3 *\*P<* 0.05. (**C&F)** Concatenate multiple flow samples to visualize CD84 and Jaml1 expression in one feature plot across all samples including; (FMO, Bone marrow, lung, spleen, MFP and tumor from WT and PyMT); significant expression was only observed in spleen and tumor from PyMT. **g**, Overview of PBMC collection, culture condition and FACS approach. (**H)** Concatenate multiple flow samples to visualize CD84 expression G- and M-MDSCs in one feature plot across all samples including PBMC control and treated. (**I-J)** Statistical analysis using unpaired t-test (Mean ± SEM of n = 3) *\*P<* 0.05

To determine how generalizable these markers are, we next explored if CD11b^+^Gr1^+^ cells express CD84 in two additional mouse models of breast cancer: a BRCA1/p53-driven model (Brca1^f11/f11^p53^f5&6/f5&6^Cre^c^)^46^ and an orthotopic transplant model using 4T1 breast cancer cells in Balb/c mice. First, we profiled the expansion of CD11b^+^Gr1^+^ cells in tumor-bearing BRCA1/p53 mice in comparison to WT mice. We observed a significant increase in CD11b^+^Gr1^+^ cells in BRCA1/p53 mice in the spleens, lungs, and tumors compared to WT (**fig. S4F**). Similar to our PyMT model, we confirmed a high proportion of CD11b^+^Gr1^+^ cells that expressed CD84 (∼24%) in spleen and (∼39.13%) in the tumor, but not in the bone marrow, or lungs, (**fig. S5A-B**). In line with these findings, we observed a significant expansion of CD11b^+^Gr1^+^ cells in the 4T1 model in bone marrow, lungs, spleens and tumors compared to WT (**fig. S4G**), and CD84 expression was elevated in spleens (∼ 21.46%) and tumors (∼ 8.49%) (**fig. S5C-D**) to a significant but lower extent compared to the other two breast cancer models (**fig. S5D**), while CD11b^+^Gr1^+^ cells from bone marrow and lungs showed no detectable CD84 expression (**fig. S5C).** Additionally, we used *in vitro* generation of MDSCs by treating myeloid cells with GMCSF^7^. We found that after G-MCSF treatment the CD11b^+^Gr1^+^ population exhibited a significant increase in CD84 positive cells (∼ 24.45%; **fig. S5E-F**) and Jaml positive cells (∼ 11.26%; **fig. S5E and G**). Finally, we used previously established protocols for *in vitro* generation of human MDSCs by isolating peripheral blood mononuclear cells (PBMCs) and treating them with G-MCSF and IL6^47^ (**Fig. 4G; fig. S4H**). We observed a significant upregulation of CD84 in samples from *in vitro* generated CD11b+/CD14+ M-MDSCs and CD11b+/CD15+ G-MDSCs compared to control cells (**Fig. 4H-J**). Together, these experiments established CD84 and Jaml as novel, generalizable cell surface markers for MDSC detection.

### CD84+ MDSCs exhibit T cell suppression and increased ROS production

To functionally validate whether CD45^+^CD11b^+^Gr1^+^CD84^hi^ cells inhibit immune cell activation, we performed co-cultures activated T cell as described above (**Fig. 5A**). Indeed, CD45^+^CD11b^+^Gr1^+^CD84^hi^ cells from spleen of tumor-bearing mice suppressed CD4 and CD8 T cell proliferation in comparison to CD45^+^CD11b^+^Gr1^+^ cells isolated from control mice (**Fig. 5B-C**). Next, we subfractionated CD11b^+^Gr1^+^CD84^low^ and CD11b^+^Gr1^+^CD84^hi^ cells from spleens and tumors of PyMT tumor-bearing mice and measured their potential for ROS production as a hallmark for MDSC function. We utilized H_2_DCFDA staining for ROS in combination with flow cytometry and observed that CD11b^+^Gr1^+^CD84^hi^ cells produced significantly higher amounts of ROS compared to CD11b^+^Gr1^+^ cells from control mice, while CD11b^+^Gr1^+^CD84^low^ showed no statistically different ROS production (**Fig. 5D-E**). Finally, we used qPCR to interrogate selected genes from our MDSCs signature and found elevated expression of the complete panel of MDSC-related genes in CD11b^+^Gr1^+^CD84^hi^ cells compared to CD11b^+^Gr1^+^CD84^low^ (**fig. S6A**). These findings indicate that MDSCs capable of T cell suppression and ROS production can be faithfully detected and enriched for based on high CD84 expression.

**Fig. 5.**
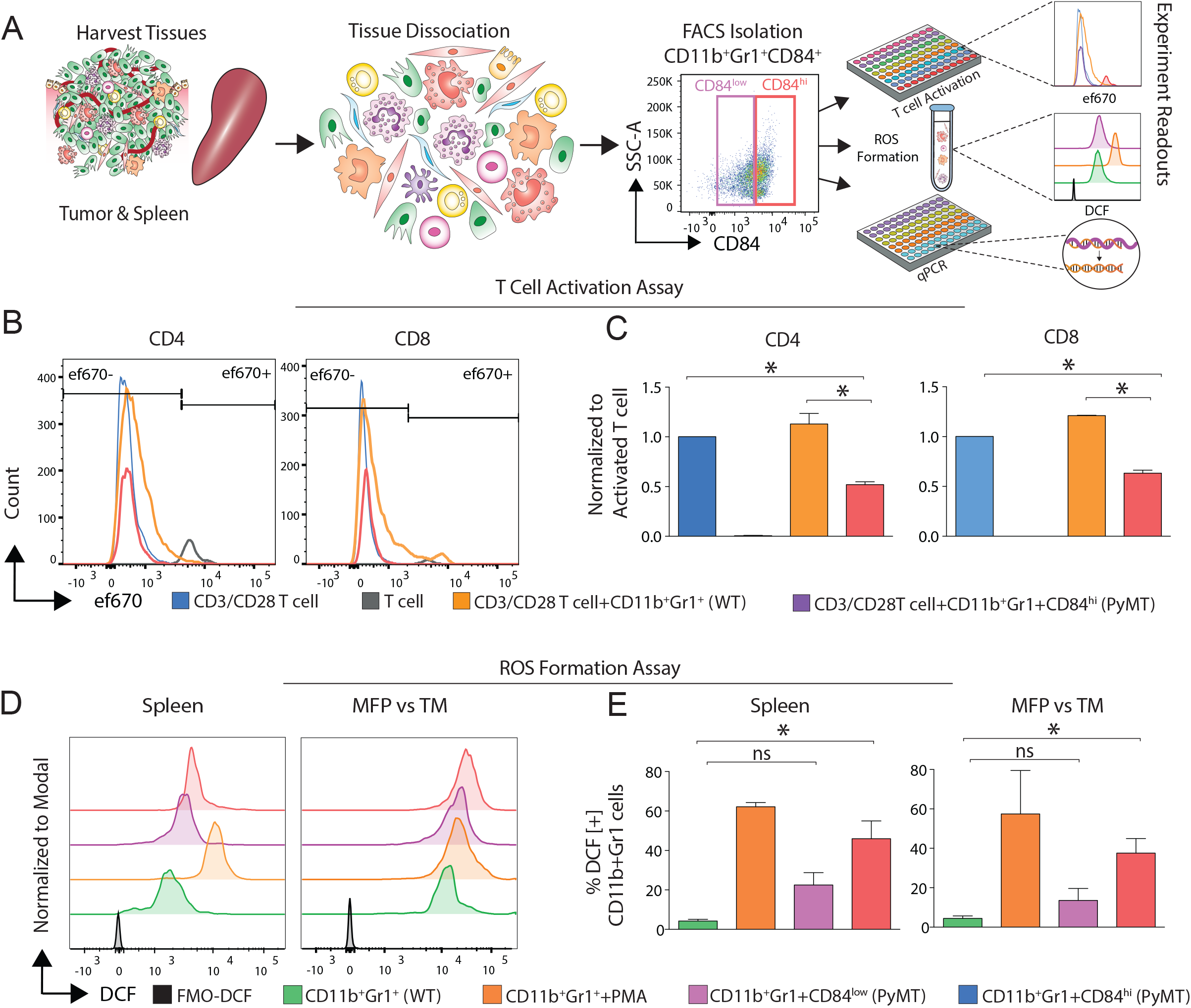
CD11b^+^Gr1^+^CD84^hi^ cells exhibit potent capacity for T cell suppression and increased ROS production. Overview of FACS approach using two different tissues (spleen and primary tumor) from WT and PyMT were subjected to T cell activation, ROS formation and qPCR assays. (**B-C)**, Splenic CD11b^+^Gr1^+^CD84^hi^ cells from tumor-bearing mice suppress T cell proliferation. Histogram overly and quantitative bar charts **(C)** showing CD4/CD8 T cell proliferation measured by FACS in control samples (T cells; red), T cells activated by CD3/CD28 (blue), activated T cells plus CD11b^+^Gr1^+^ cells from control spleens (orange) and activated T cells plus CD11b^+^Gr1^+^CD84^hi^ cells from spleen of tumor-bearing mice (purple). **c**, Statistical analysis (Mean ± SEM of n = 3) *\*P<* 0.05 One-way ANOVA. (**D-E)** CD11b^+^Gr1^+^CD84^hi^ cells from tumor-bearing mice show increased ROS formation compared to CD11b^+^Gr1^+^CD84^low^; PMA-treated cells were used as positive control. ROS was measured by FACS using H_2_DCFDA. (**E)** Statistical analysis of ROS assay unpaired t-test (Mean ± SEM of n = 3) *\*P<* 0.05.

## Discussion

Understanding the cellular and molecular mechanisms through which the tumor microenvironment can suppress an active anti-tumor immune response will be critical to improve current approaches for cancer immunotherapy such as checkpoint inhibition (*e.g.*, PD1, CTLA4) or CAR-T cell treatments^48^. MDSCs represent such microenvironmental components that commonly expand in cancer patients and promote advanced tumor progression and T cell-suppression in various different cancers including breast cancer^5^. Here, we generate the first single-cell transcriptomics map of MDSC maturation during cancer to dissect the unique molecular features of MDSCs in breast tumor-bearing mice, and to elucidate how these immunosuppressive cells differ from their normal myeloid counterparts. Using this resource, we establish an MDSC-specific gene signature that is largely shared between G- and M-MDSCs but strongly differs from their normal myeloid counterparts; we reconstruct their unique differentiation trajectory from neutrophil progenitors through an aberrant path of differentiation; and we identify novel MDSC-specific cell surface markers for detection and prospective isolation of MDSCs.

The MMTV-PyMT mouse model for breast cancer is one of the most widely used model system for studying breast cancer, which closely resembles human pathogenesis^14^ and which is known to induce significant expansion of MDSCs during tumor progression^2^. We show here that spleen is the major organ site in which MDSCs can be robustly detected. To identify unique molecular features associated with MDSC function, we utilized scRNAseq as a powerful, unbiased method to reveal hidden variation on a single-cell level in a population of FACS-isolated CD45^+^CD11b^+^Gr1^+^ cells from the spleen of WT control and tumor-bearing PyMT mice. This dataset not only provides the first single cell-level depiction of monocyte/neutrophil heterogeneity in the spleen under steady-state conditions (WT mice), but also elucidated how MDSCs emerge as distinct clusters in both monocytes and neutrophil-like cells, which allowed us to establish MDSC-specific gene signatures. Interestingly, there was significant overlap between G- and M-MDSCs, suggesting that both monocytes and neutrophils acquire similar immune-suppressive features. The MDSC signature includes various genes associated with immune regulations such as *Arg2* and *Cd84*, as well as chemokine receptors (e.g. *Ccr2*, *Cxcr2*) indicating that MDSCs are responsive to neutrophil/MDSC-recruiting chemokines guiding their migration to active sites of inflammation such as the primary tumor or metastatic foci (**Fig. 6**).

**Fig. 6.**
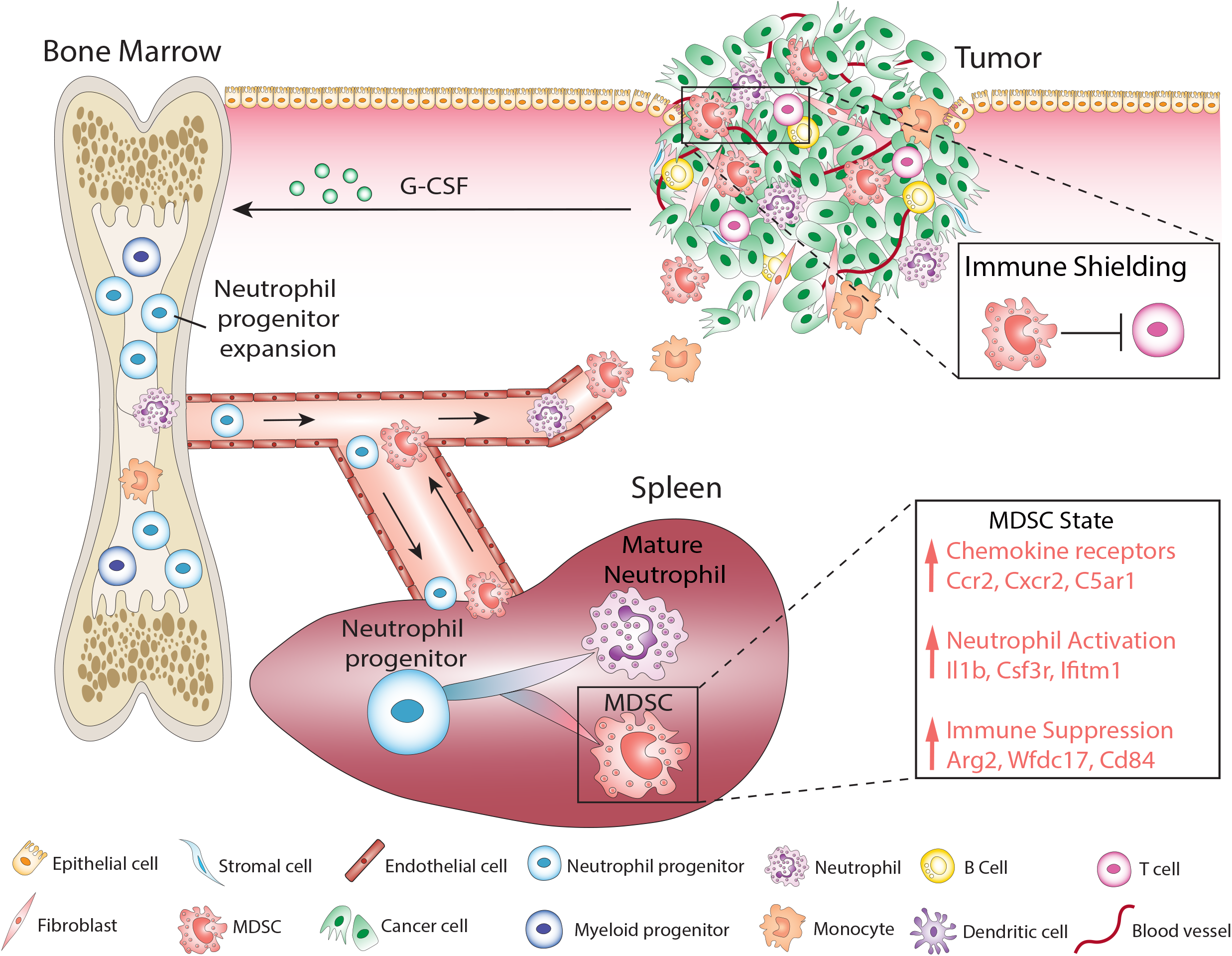
Proposed model of aberrant neutrophil differentiation in the spleen during cancer. Myeloid cells differentiate in bone marrow from hematopoietic stem cells through common myeloid progenitors. Common granulocyte/monocyte progenitors expand in the bone marrow of tumor-bearing mice and migrate to spleen as a marginated pool, where they give rise to normal neutrophil maturation and, in cancer, aberrant neutrophil differentiation into MDSCs. Our findings indicate that the MDSC-specific gene signature is largely shared between G- and M-MDSCs, but strongly differs from their normal myeloid counterparts. This MDSC signature includes immune-suppressive factors, markers of increased neutrophil activation, and numerous chemokine receptors, which likely guide their migration towards primary tumor or metastatic sites (indicated by arrows), where they may shield tumor cells from anti-tumor adaptive immunity.

Our findings indicate that G-MDSCs may emerge from neutrophil progenitors through an aberrant differentiation trajectory giving rise to a cell state that is not present in normal conditions. Interrogating our observed cell states using pseudotemporal ordering and comparing these to recent work that defined the signatures of various haematopoietic stem and progenitor cell states in single-cell resolution^17^ allowed us to reconstruct that MDSCs form an aberrant trajectory from neutrophil progenitor cells that occurs at the cost of normal differentiation into mature neutrophil granulocytes, which are less abundant in tumor-bearing mice (**Fig. 3A**). Further interrogation of the initial transitional cell state that branches off into G-MDSCs revealed several genes that strongly increase in expression in this transitional phase (**Fig. 3C**), which may suggest that therapeutically interfering with these gene products could block MDSC differentiation before they become functionally active.

A major limitation for current research studying MDSCs is the lack of specific cell surface receptors for detection and prospective isolation for functional interrogation. Here, we identify CD84 and Jaml as novel cell surface receptors on MDSCs, which can be used in combination with CD11b/Gr1 staining to detect the presence of MDSCs in various organs of tumor-bearing mice, or in human in combination with CD11b/CD14 or CD15. CD84 is involved in cell-cell interactions and modulation of the activation and differentiation of a variety of immune cells^49^, and functions as a homophilic adhesion molecule on B cells, monocytes and, on a lower extent, T cells where it enhances IFNg secretion and activation^43, 44^. Interestingly, CD84 can regulate PD-1/PD-L1 expression and function in chronic lymphocytic leukemia resulting in suppression of T cell responses and activity^50^, suggesting that CD84 may allow MDSCs to directly regulate immune checkpoints in breast cancer patients. Our future work will be focusing on functional interrogation of CD84 and other key genes identified here in MDSC biology and their capacity to inhibit T cell activation, as well as the validation of CD84 and Jaml as markers for tumor-associated MDSCs in human cancer patients.

## Materials and Methods

### Mice

All mouse experiments were approved by the Institutional Animal Care and Use Committee of University of California Irvine, in accordance with the guidelines of the National Institutes of Health. Transgenic PyMT (MMTV-PyMT) mice were purchased from The Jackson (JAX) Laboratory (stock no: 002374) and breedings were maintained on FVB/n and PyMT (MMTV-PyMT) backgrounds. Littermates from control and transgenic mice were used for all experiments.

## Tissue Collection and Cell Isolation

### Bone marrow

After mouse dissection, bone marrow (BM) was flushed from mouse tibia and femurs using a 28G needle and plastic syringe and then kept in HBSS (Corning, 21-023-CV). BM cells were centrifuged at 500g at 4°C for 5 min. Cells were incubated for 5 min at RT in 2 mL red blood cell (RBC) lysis buffer. Cells were quenched with 10 ml HBSS containing 2% FBS (Omega Scientific, FB-12) and centrifuged at 500g at 4°C for 5 min. Cells were resuspended in 3 mL FACS buffer (1xPBS, 3% FBS) and total remaining live BM cells were counted using the automated cell counter Countess™ II (ThermoFisher Scientific, AMQAX1000).

### Spleen

The spleen was pushed through a 70-μm cell strainer and washed with RPMI to create a cell suspension of splenocytes. Cells were centrifuged at 500g at 4°C for 5 min and then incubated for 5 min in 5 mL RBC lysis buffer at RT. Cells were quenched with 10 mL RPMI (Corning, 10-040-CV) with 5% FBS and centrifuged at 500g at 4°C for 5 min. Cells were resuspended in 3 mL FACS buffer (1xPBS, 3% FBS), and total remaining live cells were counted by Countess™ II and processed for FACS.

### Lung, tumor and mammary fat pad (MFP)

Tissue samples were harvested from mice and mechanically dissociated using a razor blade. Tissues were placed in DMEM/F-12 (Corning, MT10090CV) complete medium containing 5 μg/mL insulin (Sigma-Aldrich, 11376497001), 50 μg/mL penicillin/streptomycin (HyClone, SV30010), and 0.1 mg/mL collagenase type IV (Sigma-Aldrich, C5138) and were digested at 37 °C on a shaker for 45 min. Samples were centrifuged at 500g for 5 min at RT. Cells were resuspended in HBSS and centrifuged at 500g for 5 min at RT. Cells were resuspended in 25 μL DNase I (Sigma-Aldrich, D4263) for five min at RT and then 2 mL of 0.05% Trypsin (Corning, 25-052-CI) was added and samples incubated at 37°C for 10 min. Samples were centrifuged at 500g for 5 min at RT and then resuspended in 5 mL HBSS with 2% FBS. The cell suspension was filtered through a 70 μM cell strainer (Fisher Scientific, 22363548) and incubated for 5 min at RT in 3 mL RBC lysis buffer. Cells were quenched with 10 mL HBSS with 2% FBS and centrifuged at 500g for 5 min at RT. Cells were then resuspended in RPMI with 10% FBS, and total remaining live cells were counted by Countess™ II and processed for FACS.

### Peripheral blood

Blood was collected using a 20G needle and syringe from the chest cavity after the right atrium and left ventricle were punctured. Mice were perfused with 15 mL of 10 mM EDTA in 1xPBS and blood was collected. Blood cells were centrifuged at 500g at 4°C for 5 min. Cells were resuspended in 5 mL RBC lysis buffer and incubated at RT. for 5 min. Cells were quenched in 5 mL RPMI with 3% FBS and centrifuged at 500g at 4°C for 5 min. Cells were then resuspended in 3 mL FACS buffer (1xPBS, 3% FBS), and total remaining live cells were counted by Countess™ II and processed for FACS.

### Brain

Brain tissue was dissociated into a single cell suspension using the Adult Brain Dissociation Kit (Miltenyi Biotec, 130-107-677) according to the manufacturer’s protocol, and a gentleMACS Octo Dissociator with Heaters (Miltenyi Biotec, 130-096-427). Briefly, the brain was dissected from the cranium, the meninges were removed, and the brain was chopped into 8-10 pieces. The chunks were transferred into a gentleMACS C Tube (Miltenyi Biotec, 130-093-237) containing enzymes A and P, and then placed onto the gentleMACS. The brain was digested using the 37C-ADBK protocol on the instrument. After a 30-minute heated digestion, the brain slurry was strained through a 70 µm nylon strainer and washed with 10 mL ice cold 1xPBS with 2% BSA (Sigma, A-964). The suspension was centrifuged at 300g for 10 min at 4°C and then mixed with 4 mL of 1X Debris Removal Solution provided in the kit and centrifuged for 10 min at 3000g at 4°C or RT. The myelin layer was removed, and the cells were washed with DPBS, and centrifuged for 10 min at 1000g. Red blood cells were lysed for 3 min on ice with 1mL of 1X Red Blood Cell Lysis Buffer provided in the kit. After quenching in 2 mL of DPBS with 2% BSA, the cells were pelleted at 500g for 3 min at 4°C and total remaining live cells were counted by Countess™ II and processed for FACS.

## In vitro generation of MDSCs

### Mice

BM cells were collected as described above then cells were culture with RPMI and 10%FBS and treated with 20ng/ml recombinant murine (GM-CSF, Peprotech, 315-03) on day 1 and on day 3. Thus, Cells were collected on day 4 and total remaining live cells were counted by Countess™ II and processed for FACS.

### Human

We followed established protocols for *in vitro* generation of human MDSCs^47^. Briefly, Human blood were incubated with 3% dextran (Sigma-Aldrich, 31392-10G) for 18 min, supernatants were collected and followed by differential density gradient separation (Ficoll-paque^TM^ PLUS, Neta Scientific, GHC-17-1440-02). Samples were centrifuge at 500 RCF for 30 min at 20°C. PBMCs including granulocytes were collected and incubated with 10ng/ml of recombinants human cytokines (GM-CSF, Peprotech, 300-03 and IL-6, Peprotech, 200-06) or without in RPMI contain 10%FBS. Cells were treated with these cytokines every day and on day 3 cells were collected from cultures and total remaining live cells were counted by Countess™ II and processed for FACS.

### Fluorescence-Activated Cell Sorting

Tissue samples were harvested from mice and mechanically dissociated to generate single cell suspensions as described above. Cells were blocked with anti-mouse FcγR (CD16/CD32) (BioLegend, 101301) on ice for at least 10 min. Cells were then centrifuged at 500g for 5 min at 4°C and washed once with FACS buffer (1xPBS with 3%FBS). Cells were incubated for 30 min at 4°C with pre-conjugated fluorescent labeled antibodies with the following combinations: CD45 (30-F11) (BioLegend, 103112 (APC) or 103115 (APC-cy7)), CD11b (M1/70) (BioLegend, 101206 (FITC) or 101212 (APC), Gr1 (Rb6-8C5) (BioLegend, 101206 (PE) or 108439 (BV605), CD84 (mCD84.7) (BioLegend, 122805 (PE)), and Jaml (4e10) (BioLegend, 128503 (PE)). Sytox Blue dye (Life Technologies, S34857) was added to stained cells to assay for viability. Cells sorted by BD FACSAria™ Fusion and desired populations were isolated for different experiments. Human PBMCs were prepared as described above, cells were blocked with human TruStain FcX (BioLegend, 422301) on ice for 10 min. Then cells were centrifuged at 500g for 5 min at 4°C and washed once with FACS buffer (1xPBS with 3%FBS). Cells were incubated for 30 min at 4°C with the following anti-human, pre-conjugated fluorescent labeled antibodies: CD45 (efluor 450) (Thermofisher, 48-9459-42), CD11b (BV650) (BioLegend, 101206), CD14 (PerCP-Cy5.5) (Thermofisher, 45-0149-42), CD15 (APC) (BioLegend, 301907), and CD84 (PE) (BioLegend, 326007). Sytox Green (Life Technologies, S34860) was added to the stained cells to assay viability. Cells were analyzed by BD FACSAria™ Fusion.

### T cell Suppression Assay

Spleens were dissected, filtered into a single-cell suspension and depleted of red blood cells using Tris-acetic-acid-chloride (TAC). T cells were isolated from the spleen using the EasySep™ Mouse T cell Isolation Kit (StemCell Technologies, 19851) according to the manufacturer’s instructions. Isolated T cells were washed once with PBS and resuspended at 15 x10^6^/mL in staining buffer (0.01% BSA in PBS). T cells were stained with proliferation dye eFluor™ 670 (ThermoFisher Scientific, 65-0840-85) using 5mM dye per 1×10^7^ cells and incubated in a 37°C water bath for 10 min. Finally, T cells were washed and resuspended at 1×10^6^/mL in RPMI 1640 w/ HEPES+ L-glutamine (Gibco, 22400-105) complete medium containing 10% FBS (Atlanta Biologicals, S11150), 1X non-essential amino acids (Gibco, 11146-050), 100U/mL penicillin-100μg/mL streptomycin (Gibco, 15140163), 1mM sodium pyruvate (Gibco, 11360-070), and 55 μM β-mercaptoethanol (Gibco, 21985-023), eFluor™ 670-labeled T cells were plated (50×10^3^/well) in a U-bottom 96-well plate (VWR, 10062-902) and activated with plate bound anti-Armenian hamster IgG (30μg/mL, Jackson Immuno research, 127-005-099) with CD3 (0.5 μg/mL, Tonbo, 70-0031) and CD28 (1 μg/mL, Tonbo, 70-0281). Sorted CD11b^+^ Gr1^+^ cells from PyMT or WT mice were added to T cells in 1:1 ratio (50×10^3^ T cells:50×10^3^ CD11b^+^ Gr1^+^ cells). After 4 days of culture, cells were collected and blocked with anti-mouse CD16/32 (BioLegend, 101302), stained with Zombie Live/Dead Dye (BioLegend, 423105) and fluorescent-conjugated antibodies: CD4 (BioLegend, 100512; clone RM4-5), and CD8 (BioLegend, 100709; clone 53-6.7). Single-stained samples and fluorescence minus one (FMO) controls were used to establish PMT voltages, gating, and compensation parameters. Cells were processed using the BD LSR II or BD LSRFortessa™ X-20 flow cytometer and analyzed using FlowJo software v10.0.7 (Tree Star, Inc).

### ROS Production Assay

Cells were harvested from respective tissues and processed to single cell suspensions as described above. Cells were stained with CD45, CD11b, Gr1, and CD84 antibodies as described above. Following staining, cells were resuspended in FACS buffer and 10mM 2ʹ,7ʹ-Dichlorofluorescein diacetate (H2DCFDA) (Sigma-Aldrich, D6883) was added and incubated for 30 min at RT. Positive control cells were treated with 100 nM phorbol myristate acetate (PMA) (Sigma-Aldrich, P1585-1MG). Cells were then processed on the BD FACSAria™ Fusion and analyzed using FlowJo software v10.0.7 (Tree Star, Inc).

### Quantitative Real-Time PCR

CD11b^+^Gr1^+^ cells, CD11b^+^Gr1^+^CD84^hi^ cells and CD11b^+^Gr1^+^CD84^low^ cells were sorted by FACS and RNA were extracted by using Quick-RNA Microprep Kit (Zymo Research, R1050) following manufacturer’s instructions. RNA concentration and purity were measured with a Pearl nano spectrophotometer (Implen). Quantitative real-time PCR was conducted using PowerUp™ SYBR™ green master mix (Thermo Fisher Scientific, A25742) and primer sequences were found in Harvard primer bank and obtained from Integrated DNA Technologies (**Supplemental Table 9**). Gene expression was normalized to GAPDH housekeeping gene. For relative gene expression 2^negΔΔCt values were used and for statistical analysis ΔCt was used. The statistical significance of differences between groups was determined by unpaired t-test using Prism 6 (GraphPad Software, Inc).

### Single-Cell RNA Sequencing (scRNAseq)

FACS-isolated CD11b+/Gr1+ cells from the spleens of control WT (5 mice pooled) and tumor-bearing PyMT mice (3 mice pooled) were washed once in PBS with 0.04% BSA, resuspended to a concentration of approximately 1,000 cell/µL and loaded onto the 10X Genomics Chromium platform for droplet-enabled scRNAseq according to the manufacturer’s instructions. Library generation was performed following the Chromium Single Cell 3ʹ Reagents Kits v2 User Guide: CG00052 Rev B. Each library was sequenced on the Illumina HiSeq 4000 platform to achieve an average of 48,488 reads per cell. Alignment of 3’ end counting libraries from scRNAseq analyses was completed utilizing 10× Genomics Cell Ranger 2.1.0. Each library was aligned to an indexed mm10 genome using Cell Ranger Count. “Cell Ranger Aggr” function was used to normalize the number of confidently mapped reads per cell across the two libraries.

### Cluster Identification Using Seurat

The Seurat pipeline (version 2.3.1) was used for cluster identification in scRNAseq datasets. Data was read into R (version 3.5.0) as a counts matrix and scaled by a size factor of 10,000 and log transformed. We set gene expression cut-offs at minimum of 500 and a maximum cut-off of 5000 genes per cell for each dataset. In addition, cells with a percentage of total reads that aligned to the mitochondrial genome (referred to as percent mito) greater than 8% were removed. Using Seurat’s Canonical Correlation Analysis (CCA), cells from WT and PyMT mice were integrated together into a single analysis. For tSNE projection and clustering analysis, we used the first 20 principal components. Specific markers for each cluster identified by Seurat were determined using the “FindAllMarkers” function. For gene scoring analysis, we compared gene signatures and pathways in subpopulations using Seurat’s “AddModuleScore” function. For cell type subset analyses (Monocytes and Neutrophils), clusters with high expression of cell type markers (*Csf1r* and *Ly6g,* respectively) were subset out and standard Seurat workflow was applied on each. In the case of the neutrophil-specific analysis, a population of cells that grouped together and expressed a set of markers associated with neutrophil progenitors^17^ was manually labeled and treated as a distinct cluster for analysis.

### Reconstruction of Differentiation Trajectories using Monocle

Using the R package Monocle (version 2.8.0), a differentiation hierarchy within the neutrophil compartment was reconstructed. Starting with all cells from the WT and PyMT combined analysis, neutrophils were specifically subset out. Once subset, contaminating cell types were removed and the cells were re-clustered to explore additional heterogeneity within the neutrophils compartment. Using marker genes of these clusters, the top 20 unique genes per cluster were used to order cells along a pseudotemporal trajectory. Because cells that expressed markers associated with neutrophil progenitors^17^ localized to a single branch, that branch was chosen as the start of pseudotime for further analysis.

### Statistical Analysis

All data are expressed as mean ± SEM or SD and performed using Prism 6 software (GraphPad Software, Inc). P values were considered to be significant when p<0.05.

### Data Availability

Data will be publicly available on GEO (Accession numbers pending).

## Acknowledgements

We thank Dr. Devon Lawson for the constructive feedback on experimental design and analysis. We thank Ryan T. Davis and Linzi Megan Hosohama for careful review and feedback of the Manuscript. We thank Dr. Jennifer Atwood for assistance in BD FACSAria™ Fusion in the flow cytometry core at the University of California, Irvine. This study was supported by funding from NIH/NCI (1R01CA234496; 4R00CA181490 to K.K.), the American Cancer Society (132551-RSG-18-194-01-DDC to K.K.), the University of Hail, Hail, Saudi Arabia for the Ph.D. Fellowship (to H.A), the Canadian Institutes of Health Research (CIHR) Postdoctoral Fellowship (to D.M.), and pilot funds from the U54 Center for Cancer Systems Biology (CA217378).

## Author Contributions

H.A., L.M., D.M., Q.N., K.N., J.R., G.H., K.E., L.T., and A.S. performed research. K.K. supervised research. C.W. contributed new reagents and analytic tools. N.P., H.A., and K.K. performed bioinformatic analyses. H.A. and K.K. wrote the paper manuscript, and all authors discussed the results and provided comments and feedback.

---

**Fig. S1.**
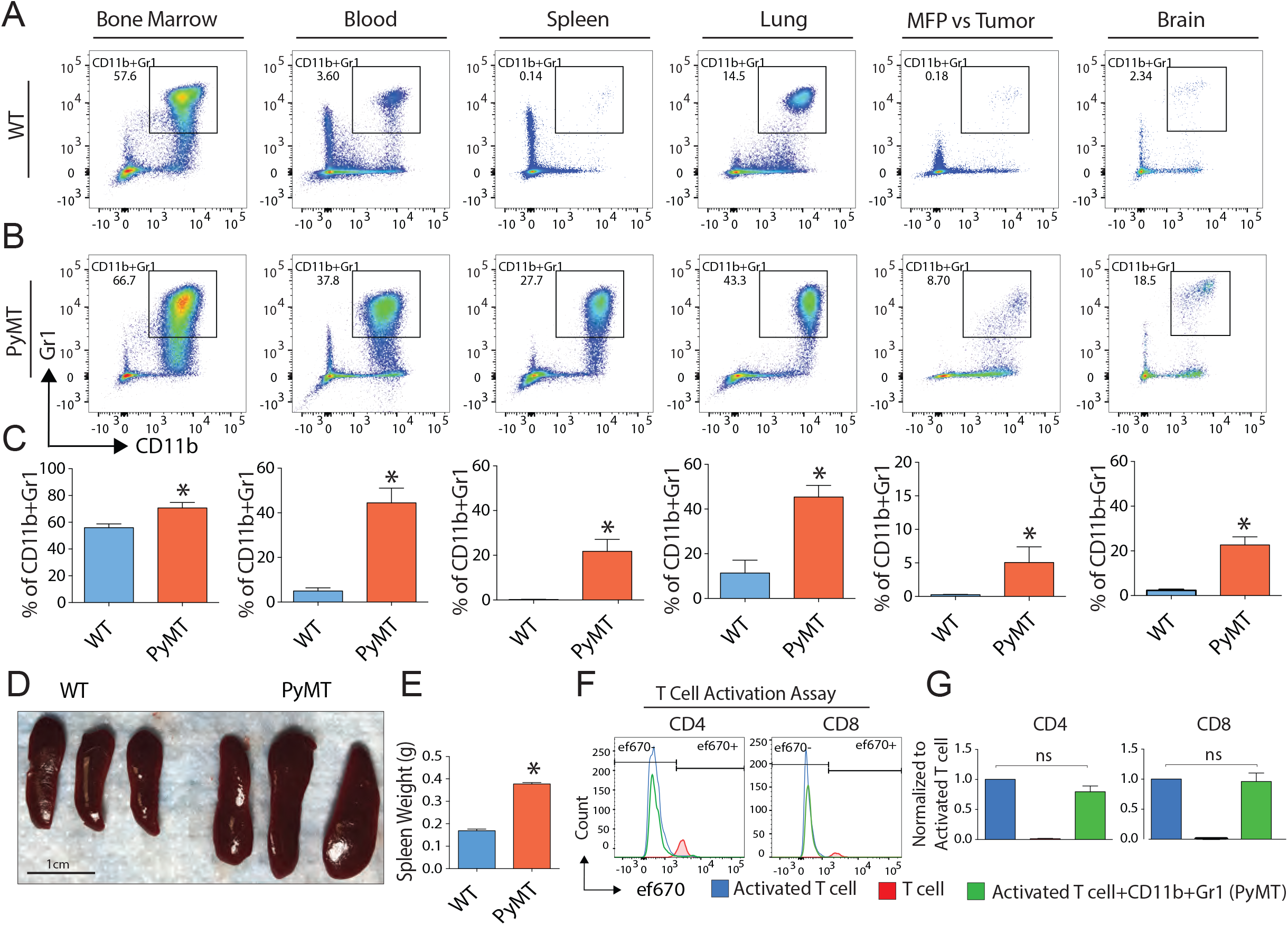
Expansion of CD11b^+^Gr1^+^ cells during tumor progression in PyMT mice. (**A-B**), Tissues from tumor-bearing PyMT and WT mice were collected and analyzed by FACS. Cells from WT **(A)** and PyMT **(B)** mice were gated on CD45+ and analyzed using CD11b/Gr1 to identify neutrophils/monocytes, which expanded significantly during tumor progression in bone marrow, blood, spleen, lung, brain and tumor compared to WT. (**C)** Combined quantification of FACS results including statistical analysis is shown in bar graphs (Mean ± SEM of n =3) *\*P<* 0.05 t-test vs. WT. **(D)** Spleen from tumor-bearing PyMT (14 weeks) was enlarged compared to WT. (**E-F**) T cell suppression assay, CD11b^+^Gr1^+^ cells were sorted from PyMT-lung and co-cluttered with activated T cells showed no effect in T cell proliferation. Statistical analysis one-way ANOVA (Mean ± SEM of n = 3) *\*P<* 0.05.

**fig. S2.**
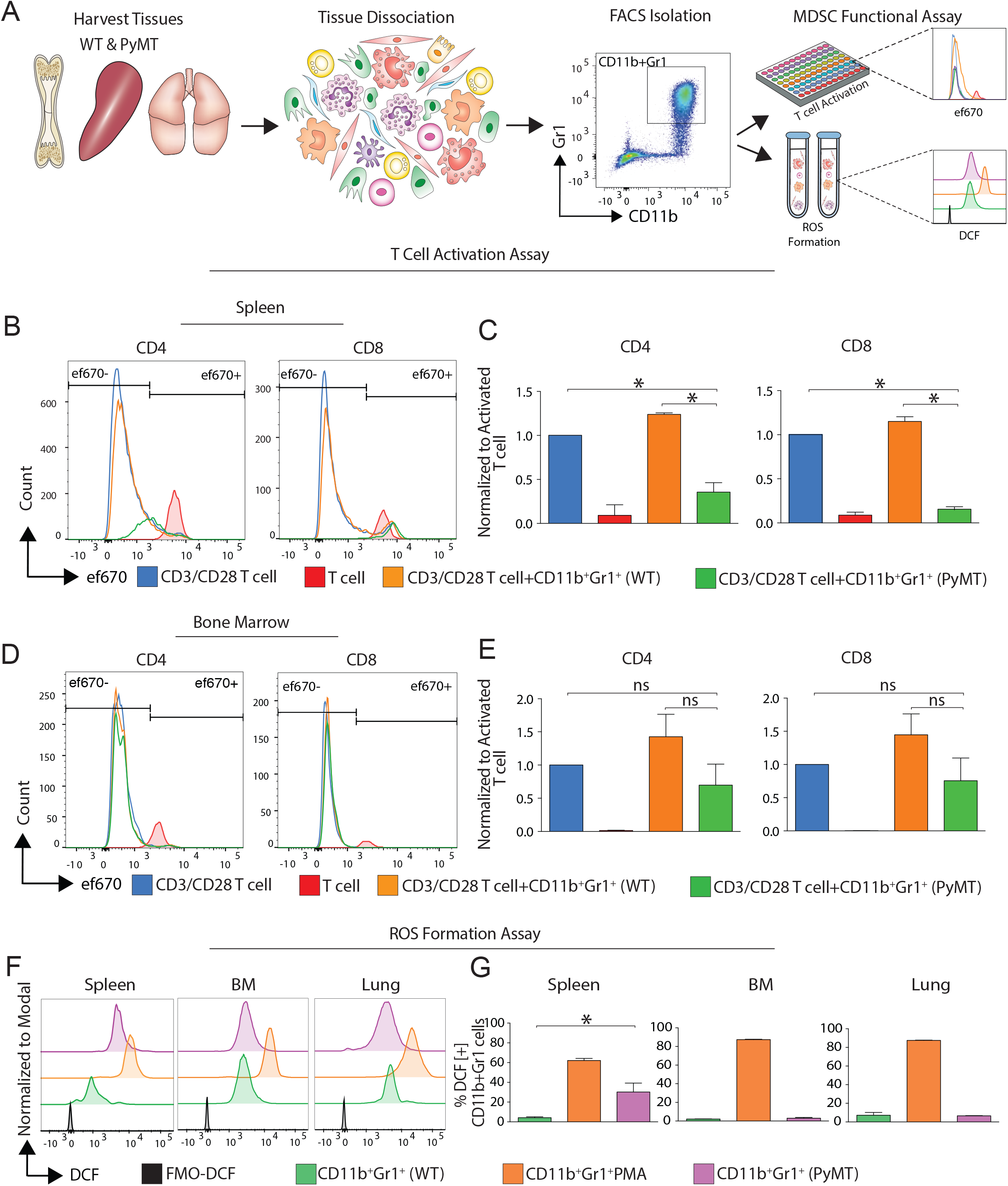
MDSCs emerge predominantly in spleen of tumor-bearing mice. (**A**) Experimental overview. Bone marrow, spleen and lung were processed into single cell suspensions and (sytox blue-negative) CD45^+^CD11b^+^Gr1^+^ cells were sorted and subjected to functional T cell suppression and ROS formation assays. (**B-C),** Splenic CD11b^+^Gr1^+^ cells from tumor-bearing mice suppress T cell proliferation. Histogram overlay **(B)** and quantitative bar charts **(C)** showing CD4/CD8 T cell proliferation measured by FACS in control samples (T cells; red), T cells activated by CD3/CD28 (blue), activated T cells plus CD11b^+^Gr1^+^ cells from control spleens (orange) and activated T cells plus CD11b^+^Gr1^+^ cells from spleen of tumor-bearing mice (green). (**D-E),** Bone marrow-derived CD11b^+^Gr1^+^ cells from tumor-bearing mice show non-significant suppression of T cell activation. Histogram overly **(B)** and quantitative bar charts (c) showing CD4 and CD8 T cell proliferation measured by FACS in T cell control samples (red), CD3/CD28 activated T cells by CD3/CD28 (blue), activated T cells plus CD11b^+^Gr1^+^ cells from control bone marrow (orange) and activated T cells plus CD11b^+^Gr1^+^ cells from bone marrow of tumor-bearing mice (green). ns = not significant. **(F-G),** CD11b^+^Gr1^+^ cells from tumor-bearing mice show increased ROS formation. PMA-treated cells were used as positive control. ROS was measured by FACS using H_2_DCFDA in CD11b^+^Gr1^+^ cells from bone marrow, spleen, and lung from control and tumor-bearing mice. Only CD11b^+^Gr1^+^ cells (MDSCs) from PyMT’s spleen significantly produced more ROS. (**C)** Statistical analysis (Mean ± SEM of n = 4) *\*P<* 0.05 One-way ANOVA. (**E)** Statistical analysis (Mean ± SEM of n = 3) *\*P<* 0.05 One-way ANOVA. (**G)** Statistical analysis unpaired t-test (Mean ± SEM of n = 3) *\*P<* 0.05.

**fig. S3.**
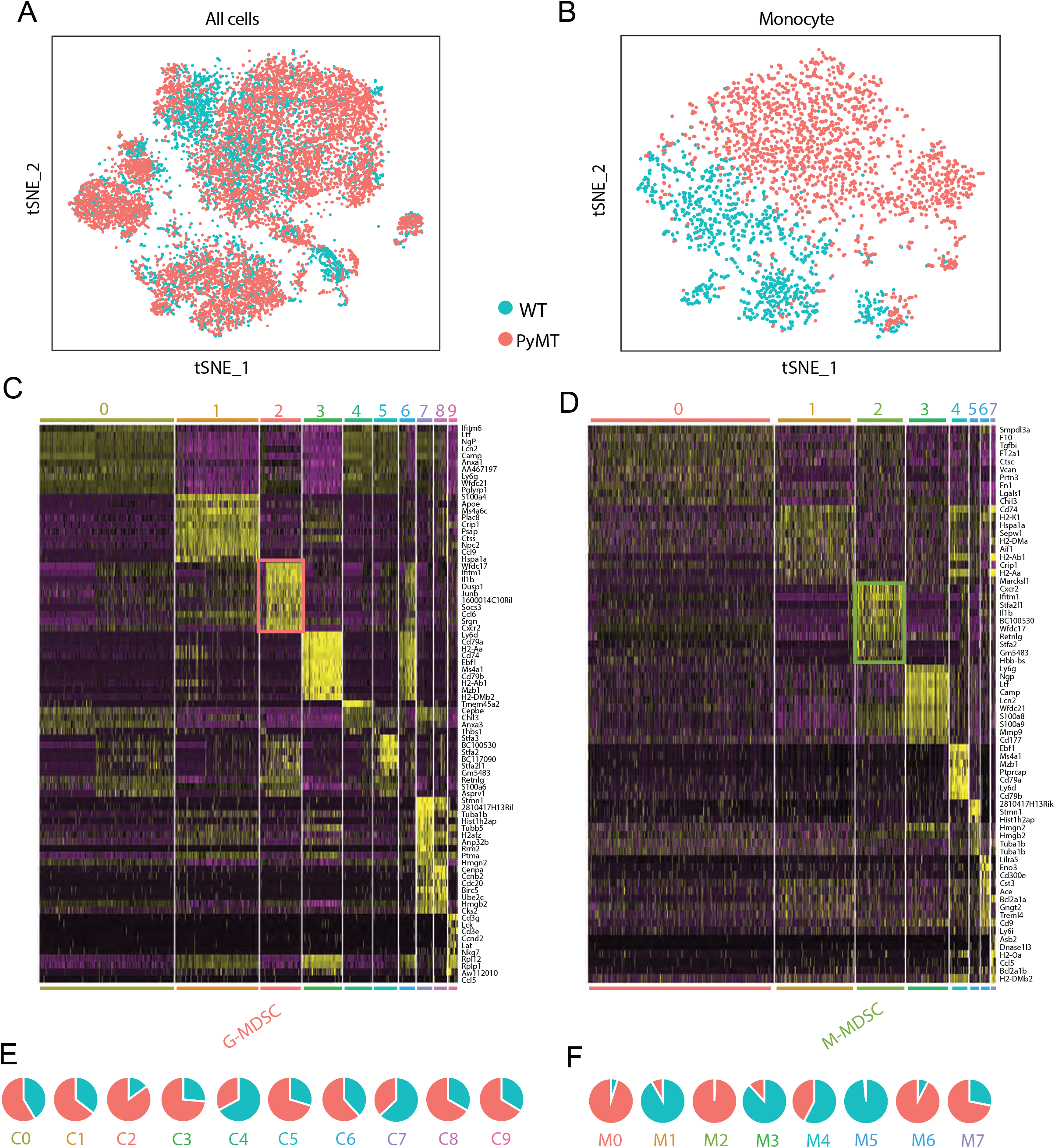
Sample labels and marker genes from scRNAseq analysis. (**A**) Seurat analysis of combined CD11b^+^Gr1^+^ cells from WT and tumor-bearing PyMT mouse spleens shown in tSNE projection labeled by tissue source (WT = blue; PyMT = red). **(B)** Seurat analysis subsetted on monocytes only from WT and tumor-bearing PyMT mouse spleens shown in tSNE projection labeled by tissue source (WT = blue; PyMT = red). (**C-D)** Heatmaps of Top 10 upregulated genes of all clusters in of combined Seurat analysis including G-MDSC cluster C2 (**C**) and subset monocyte analysis (**D**) in all monocyte clusters including M-MDSC cluster M2. (**E-F)**, Membership pie charts demonstrated clusters belong to WT or PyMT.

**fig. S4.**
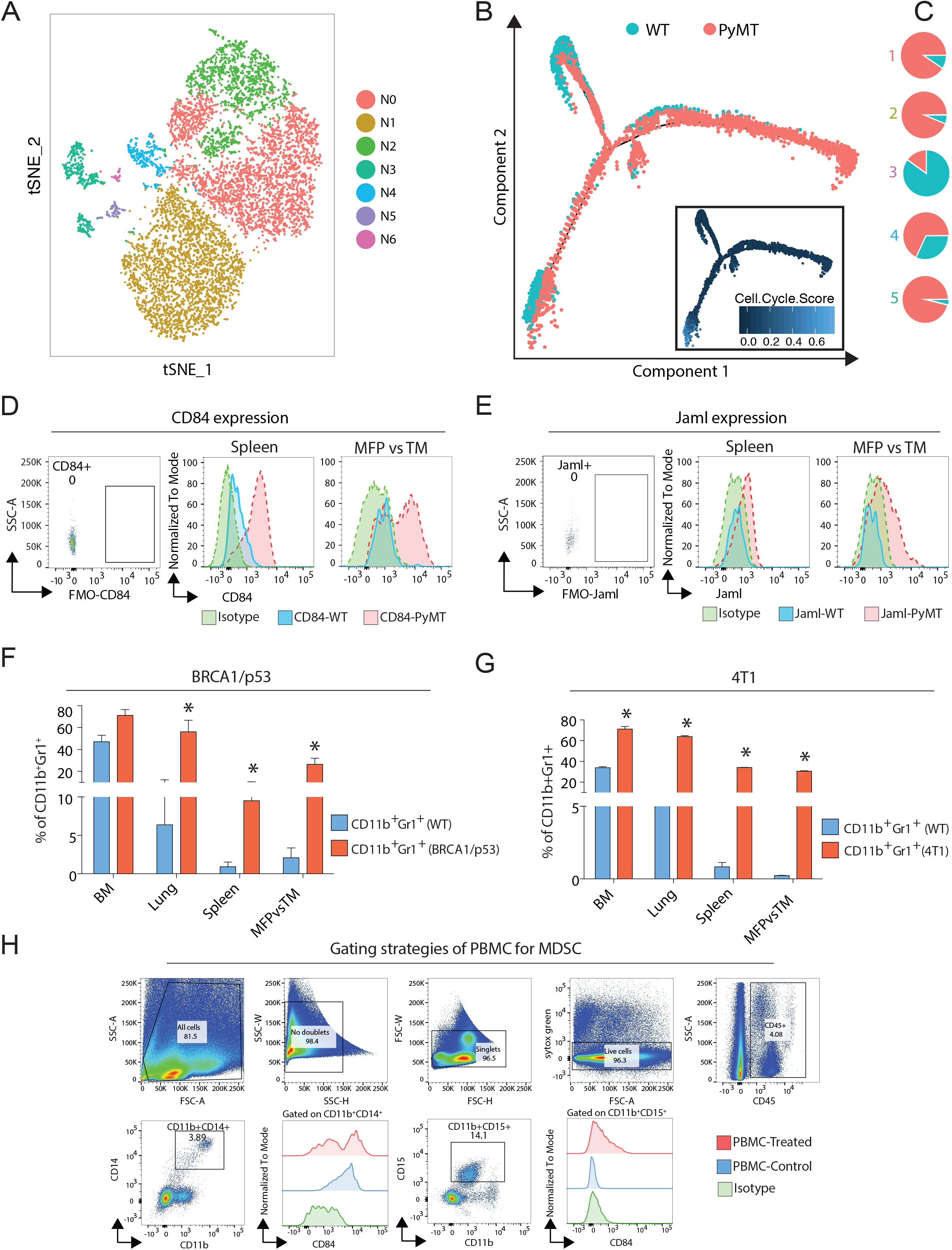
Identification of differentiation trajectory and cell surface marker expression. (**A**) Seurat-based clustering of neutrophil subset is shown, which was used define a set of marker genes for subsequent Monocle analysis. (**B**) Pseudotemporal analysis using Monocle labeled by cell source (WT=blue; PyMT=red) and cell cycle score overlay in monocle plot. (**C)** Membership pie charts per monocle detected state (WT=blue; PyMT=red). (**D-E)**, FMO and isotype controls were used to determine CD84 and Jaml expression. (**F-G)** Tissues from BCRA1/TP53 or 4T1 breast cancer models and WT mice were collected and processed to single cell suspensions. Cells were stained with antibodies for CD45^+^CD11b^+^Gr1^+^ and were gated based on live and FMO controls then analyzed by flow cytometry. (**F)** CD11b^+^Gr1^+^ cells were profiled in different tissues from WT and BRCA1/TP53 showed increase expansion significantly in tumor bearing host in spleen, lung, and tumor compared to WT. (**G)** CD11b^+^Gr1^+^ cells were profiled in different tissues from WT and 4T1 and showed increase expansion in tumor bearing host in bone marrow, spleen, lung, and tumor compared to WT. Statistical analysis unpaired t-test (Mean ± SD of n =3) *\*P<* 0.05 t-test. (**H)** gating strategies and isotype control were used (CD45^+^CD11b^+^CD14^+^ or CD15^+^) to determine CD84 expression in human G- and M-MDSCs.

**fig. S5.**
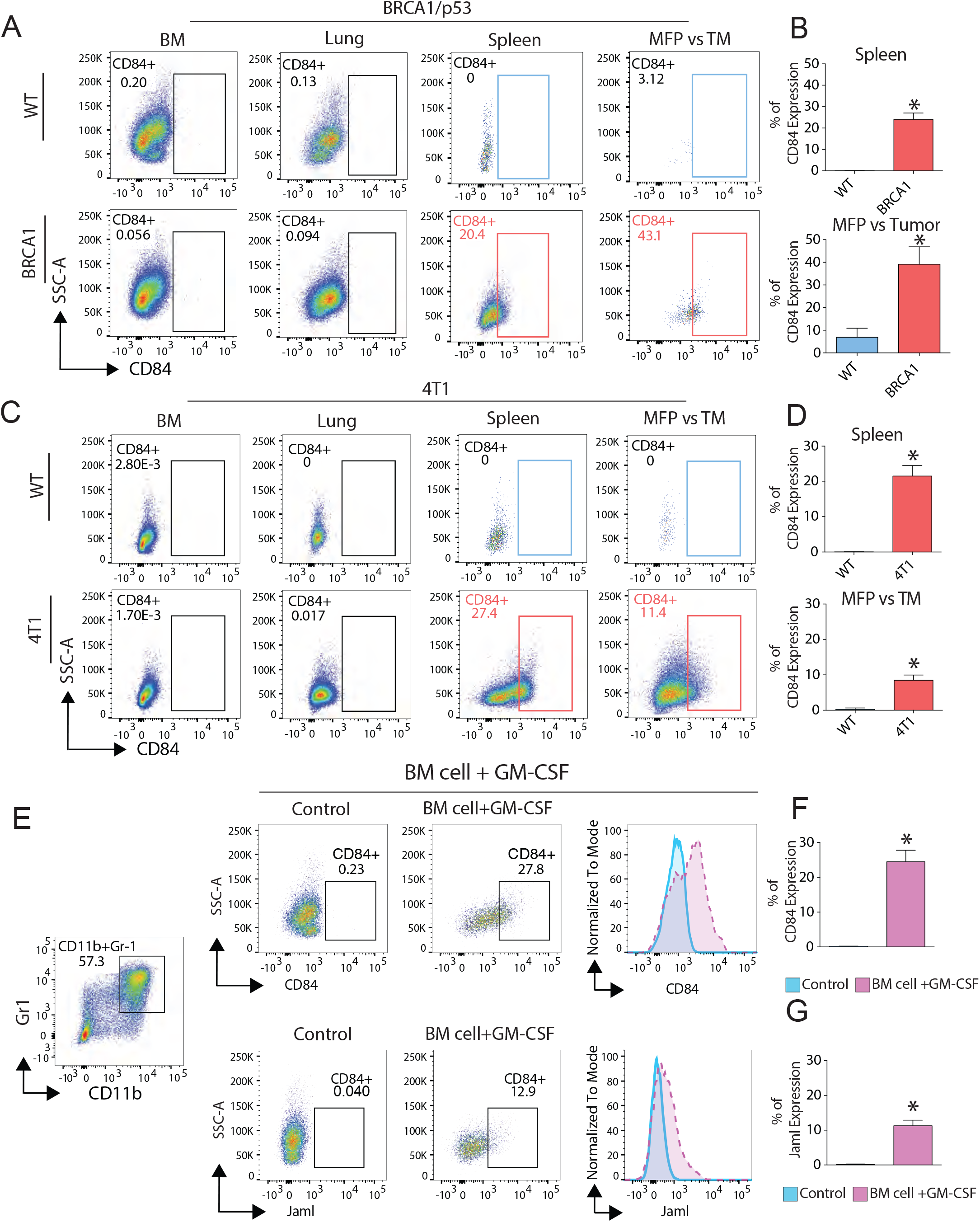
CD84 is a generalizable MDSC marker in different breast cancer models. (**A-B**) FACS plots show profiling of CD84 expression in the CD11b^+^Gr1^+^ cell population in WT and BRCA1/p53 mice. Bar charts show quantitative analysis of FACS results (Mean ± SEM of n =3). *\*P<* 0.05 **(C-D)**, FACS plots show CD84 expression in orthotopic 4T1 breast cancer model specifically in spleen and tumor compared to WT mice. Bar charts show quantitative analysis of FACS results (Mean ± SEM of n =3). *\*P<* 0.05 (**E-F)**, *In vitro* MDSCs generation by treating bone marrow cells with GM-CSF showed increase expression of CD84 and Jamal compared to untreated group normal bone marrow cells. Statistical analysis unpaired t-test (Mean ± SEM of n =3) *\*P<* 0.05

**fig. S6.**
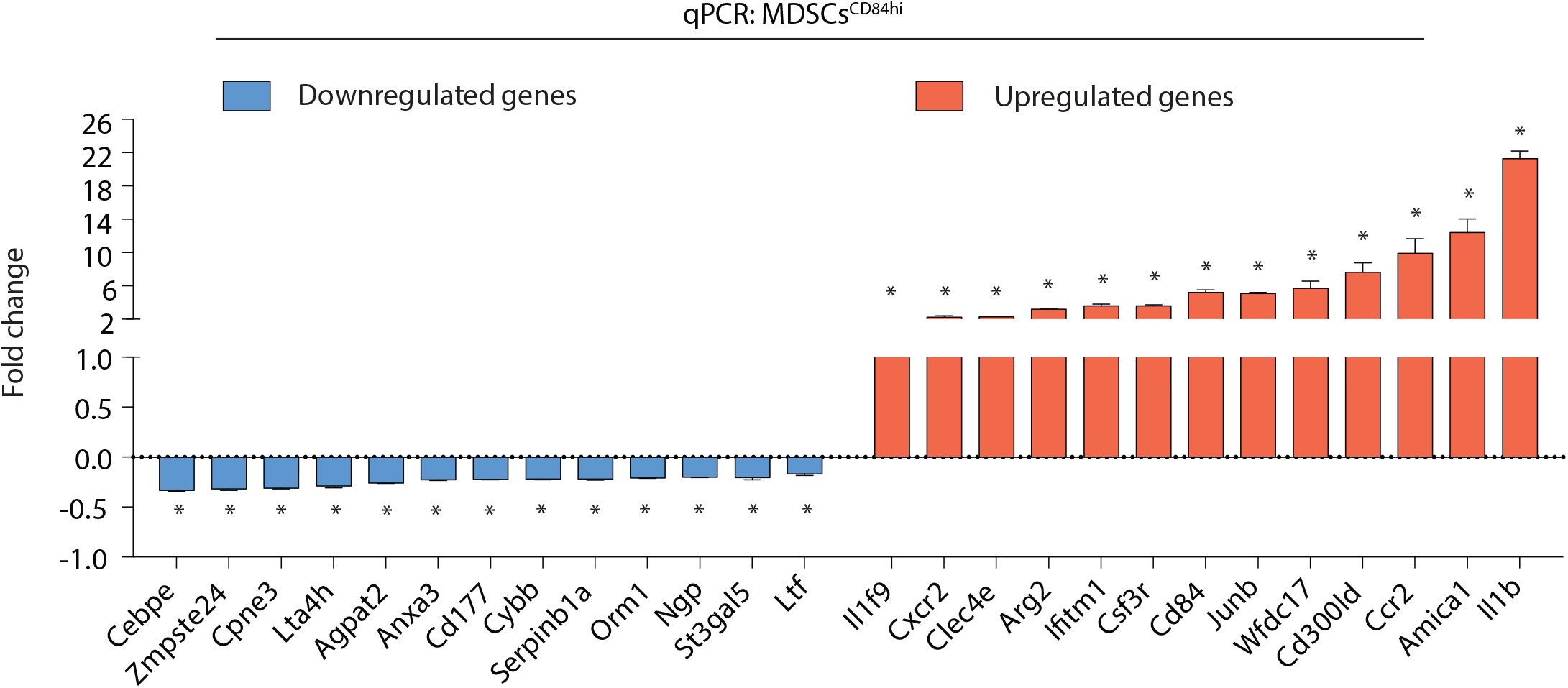
Characterization and validation of CD11b^+^Gr1^+^CD84^hi^ cells using qPCR and FACS. (**A**) CD11b^+^Gr1^+^CD84^hi^ cells and CD11b^+^Gr1^+^CD84^low^ cells were sorted by FACS and subjected to qPCR. Numerous of genes were confirmed to be significantly upregulated or downregulated in CD11b^+^Gr1^+^CD84^hi^ compared to CD11b^+^Gr1^+^CD84^low^. Statistical analysis unpaired t-test (Mean ± SEM of n =3) *\*P<* 0.05.

## SUPPLEMENTARY TABLES

**Table S1.** Marker genes from combined Seurat analysis

**Table S2.** Marker genes from Seurat analysis of monocytes only

**Table S3.** Gene signature from G-MDSCs vs. Neutrophils comparison

**Table S4.** Gene signature from M-MDSCs vs. Monocytes comparison

**Table S5.** Combined MDSC signature gene list

**Table S6.** GO terms (Biological Process 2018) MDSC gene signature

**Table S7.** Marker genes from Seurat analysis of neutrophils only

**Table S8.** Monocle state marker genes

**Table S9.** qPCR primer sequences

